# Accurate MHC Motif Deconvolution of Immunopeptidomics Data Reveals a Significant Contribution of DRB3, 4 and 5 to the Total DR Immunopeptidome

**DOI:** 10.1101/2021.11.23.469647

**Authors:** Saghar Kaabinejadian, Carolina Barra, Bruno Alvarez, Hooman Yari, William H Hildebrand, Morten Nielsen

## Abstract

Mass spectrometry (MS) based immunopeptidomics is used in several biomedical applications including neo-epitope discovery in oncology and next-generation vaccine development. Immunopeptidome data are highly complex given the expression of multiple HLA alleles on the cell membrane and presence of co-immunoprecipitated contaminants. The absence of tools that accurately deal with these challenges is currently a major bottleneck for the large-scale application of this technique. Here, we present the MHCMotifDecon that benefits from state-of-the-art HLA class-I and class-II predictions to accurately deconvolute immunopeptidome datasets and assign individual ligands to the most likely HLA allele while discarding co-purified contaminants. We have benchmarked the tool against other state-of-the-art methods and illustrated its application on experimental datasets for HLA-DR demonstrating a previously underappreciated role for HLA-DRB3/4/5 molecules in defining HLA class II immune repertoires. With its ease of use MHCMotifDecon can efficiently guide interpretation of immunopeptidome datasets, serving the discovery of novel T cell targets.

## Introduction

Mass spectrometry (MS) based immunopeptidomics is becoming increasingly relevant for several biomedical applications including cancer neoantigen discovery, vaccine development and protein drug deimmunization pipelines (Barra et al., 2020; Bettencourt et al., 2020; Mayer and Impens, 2021; Nelde et al., 2021). The broad interest in MHC-derived MS relies on its ability to discover MHC presented peptides in real scenarios, such as biomarker discovery in cancer, autoimmune and infectious disease, and therefore identifying peptides that might induce T cell responses. Additionally, thousands of peptides derived from the self-proteins can be collected in a single assay which can be directly loaded into bioinformatic pipelines to help uncovering the rules governing the processing and presentation of peptides in a biological context. Furthermore, ongoing refinements of the MS experimental workflows, such as *de novo* peptide annotation, or peptide quantification hold great promise in expanding the usage of the technique.

Despite the great potential of MS immunopeptidomics data, several inherent challenges curb the broad benefit of the technique. A major challenge being assigning the accurate MHC allele to each presented MS-derived peptide. To tackle this concern, several experimental approaches that artificially engineer cells to only express one individual HLA have been performed both for MHC class I (Abelin et al., 2017; Sarkizova et al., 2020), and MHC class II (Abelin et al., 2019). Although this laborious experimental setup has proved extremely useful to understand the rules of HLA presentation and their motifs, it lacks the biological relevance, limiting its application in other scenarios and experimental settings that involve patient samples and biospecimens.

As an alternative to experimental approaches, unsupervised bioinformatic methods have been proposed to deconvolute the peptide specificities present in complex immunopeptidomes, such as GibbsCluster (Andreatta et al., 2017; Andreatta et al., 2013) or MixMHCp (Racle et al., 2019). These pioneering tools have proved to be highly useful for MS data analysis (Alvarez et al., 2018; Mommen et al., 2016; Parker et al., 2021; Sofron et al., 2016). However, they also show intrinsic limitations such as the need for manual inspection associating those motifs to known allele specificities present in the sample and assigning peptides to MHC molecules with overlapping motifs. Another challenge for these methods is dealing with proper deconvolution of MHC molecules with limited peptide repertoire such as HLA-C. Furthermore, in MHC class II, the expression of variable alpha and beta genes forming the HLA-DQ and HLA-DP heterodimers, plus the expression of HLA-DRB1 and its associated HLA-DRB3, 4 and 5 alleles, make up to twelve potential HLA class II specificities, disputing the capacity of these methods to learn from this complex data.

Recently, Alvarez et al. proposed the NNAlign_MA framework that combines binding affinity and MS data to learn and deconvolute the peptide specificity associated to MS immunopeptidome samples on the fly, while training a neural network to predict MHC presentation (Alvarez et al., 2019). This method showed a high deconvolution accuracy, and this core algorithm has been already introduced into the state-of-the-art predictors NetMHCpan-4.1 and NetMHCIIpan-4.0 (Reynisson et al., 2020a).

Here, we build on these achievements to develop MHCMotifDecon, a user-friendly supervised tool, that benefits from the prior knowledge of NetMHCpan and NetMHCIIpan to perform MHC motif deconvolution of both class I and class II MS immunopeptidomes, assigning peptide sequences to their most likely HLA restriction. The output from the tool includes HLA motifs, length distributions, peptide counts, and a trash cluster to discard non-specific or contaminating coimmunoprecipitated peptides. To show its power in analyzing immunopeptidomics data, the tool is benchmarked against other state-of-the-art publicly available motif deconvolution methods on artificial MS datasets.

To further demonstrate the power of MHCMotifDecon, we additionally generated and motif deconvoluted novel HLA-DR immunopeptidome datasets. Considering that the DR immunopeptidome of different antigen presenting cells is naturally a complex mixture of peptides presented by HLA-DRB1 (primary DRB) and DRB3, 4 and 5 (secondary DRB) alleles, besides the distinct roles these alleles play in disease susceptibility or protection, the deconvolution of the peptide repertoire of DRB1 molecules from the DRB3, 4 and 5 is essential for unraveling the function of class II HLA-DR in the course of autoimmune disorder progression and treatment. However, due to their strong linkage disequilibrium (LD) with the accompanying DRB1 alleles, the contribution of the secondary DRB alleles in presenting antigenic peptides to T cells and their role in immune activation and response has been largely overlooked.

Here, we demonstrate how MHCMotifDecon offers a simple yet highly powerful tool to address this issue, allowing for quantification of the relative contribution of DRB3, 4 and 5 molecules to the full HLA-DR peptide repertoire.

## Results

The proposed MHCMotifDecon method allows for HLA motif deconvolution of immunopeptidome datasets based on prediction of peptide HLA restrictions. The tool takes one or multiple immunopeptidome datasets as input together with information of the HLA molecules expressed in each individual cell line/sample. In the following sections the performance of the tool is first benchmarked against other publicly available HLA motif deconvolution methods, and next the use and value of the tool is illustrated in a series of real-life applications.

### Performance evaluation

To assess the deconvolution accuracy and value of MHCMotifDecon, the method was tested and compared to other publicly available tools on two artificial datasets (one for HLA class I and another for HLA class II) constructed combining MS peptides from experimental designs where the restriction allele was known and unique. As MHCMotifDecon is a supervised classification method, the two datasets were constructed from peptides that do not share overlap with the MHC data used for the training of NetMHCpan and NetMHCIIpan (for details refer to materials and methods).

### MHC class I deconvolution

For MHC class I, 4400 HLA ligands of length 8-14 amino acids were randomly sampled from six single-allele datasets (HLA-A*02:02, HLA-A*11:02, HLA-B*13:01, HLA-B*49:01, HLA-C*07:02, HLA-C*14:03) (Abelin et al., 2017) and merged into one artificial dataset. HLA-A and HLA-B ligands were added in equal proportions and five times more compared to HLA-C (Apps et al., 2015), mimicking the biological setting where HLA-A and HLA-B alleles are expressed at higher levels compared to HLA-C.

Next, MHCMotifDecon, MixMHCp (Bassani-Sternberg and Gfeller, 2016) and GibbsCluster (Andreatta et al., 2017) were used to deconvolute the motifs contained within this artificial MHC class I dataset of 4400 ligands covering 6 HLA molecules. The result of this deconvolution is shown in figure 1. As an additional comparison of the deconvoluted motifs, each of the original single allele data was submitted to GibbsCluster (Alvarez et al., 2018) with a fixed cluster number of 1 to show the motifs found in the original MS samples (figure 1).

**Figure 1.**
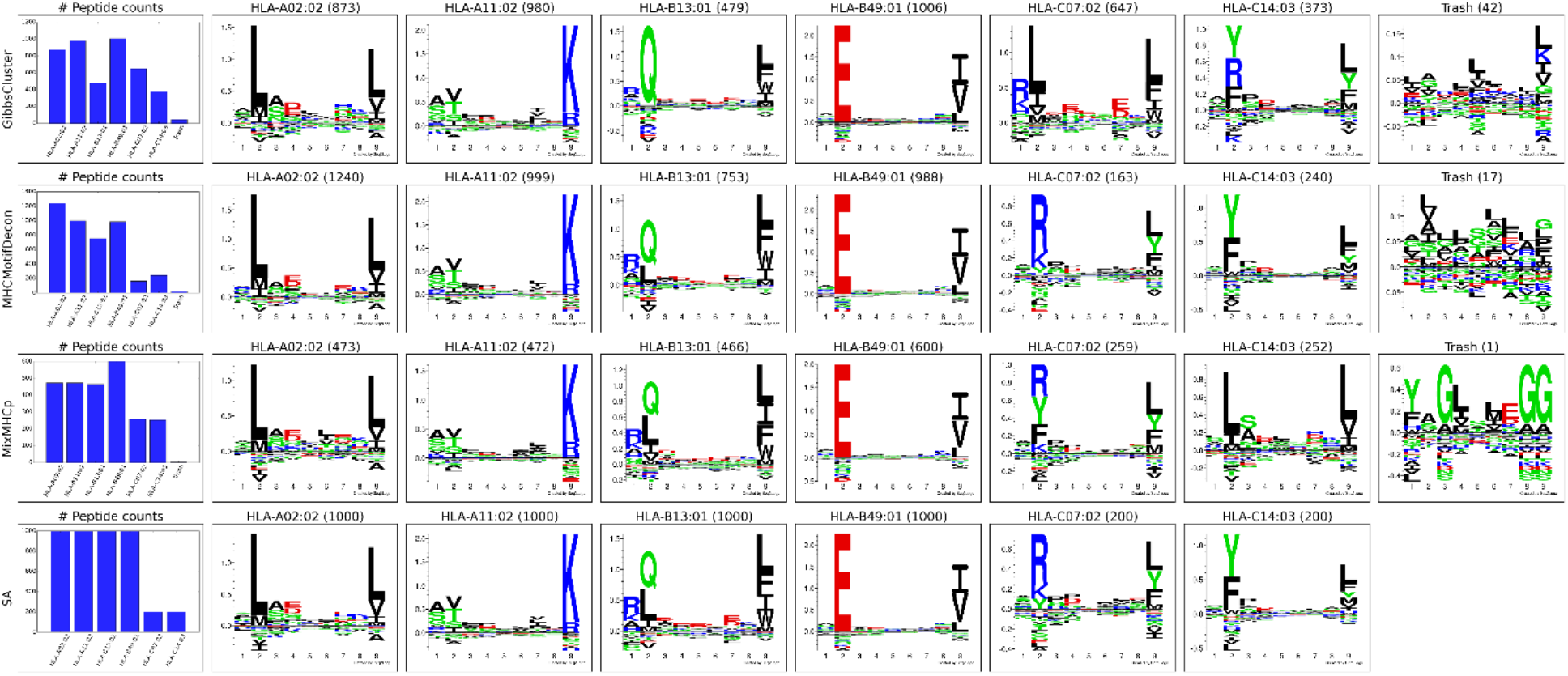
Motif deconvolution of the artificial HLA class I dataset. A. GibbsCluster, MHCMotifDecon, MixMHCp deconvolution of datasets. Motifs for MixMHCp were constructed from the set of deconvoluted 9mer peptides only. The lower panel (SA) shows motifs for each individual HLA-A*02:02, HLA-A*11:02, HLA-B*13:01, HLA-B*49:01, HLA-C*07:02, HLA-C*14:03 dataset as obtained by GibbsCluster using a single cluster.

While all clustering algorithms were able to separate most HLA-A and HLA-B associated peptides, the low representation of HLA-C peptides in combination with a less defined motif makes this motif more difficult to establish (figure 1). Focusing on the identified motifs for HLA-C*07:02 and HLA-C*14:03, it becomes further apparent that both MixMHCp and GibbsCluster fail to accurately separate the correct motifs for the two HLA-C molecules (figure 1, SA panel).

The accuracy of the motif deconvolution was next quantified in terms of a confusion matrix aligning the annotated and predicted HLA restrictions for each of the 4400 peptides in the dataset (figure 2). These results demonstrate an overall high ability of the three methods to deconvolute the motifs from HLA-A and HLA-B, but also illustrates the difficulty of the two unsupervised methods to accurately capture the motifs of the two HLA-C molecules.

**Figure 2.**
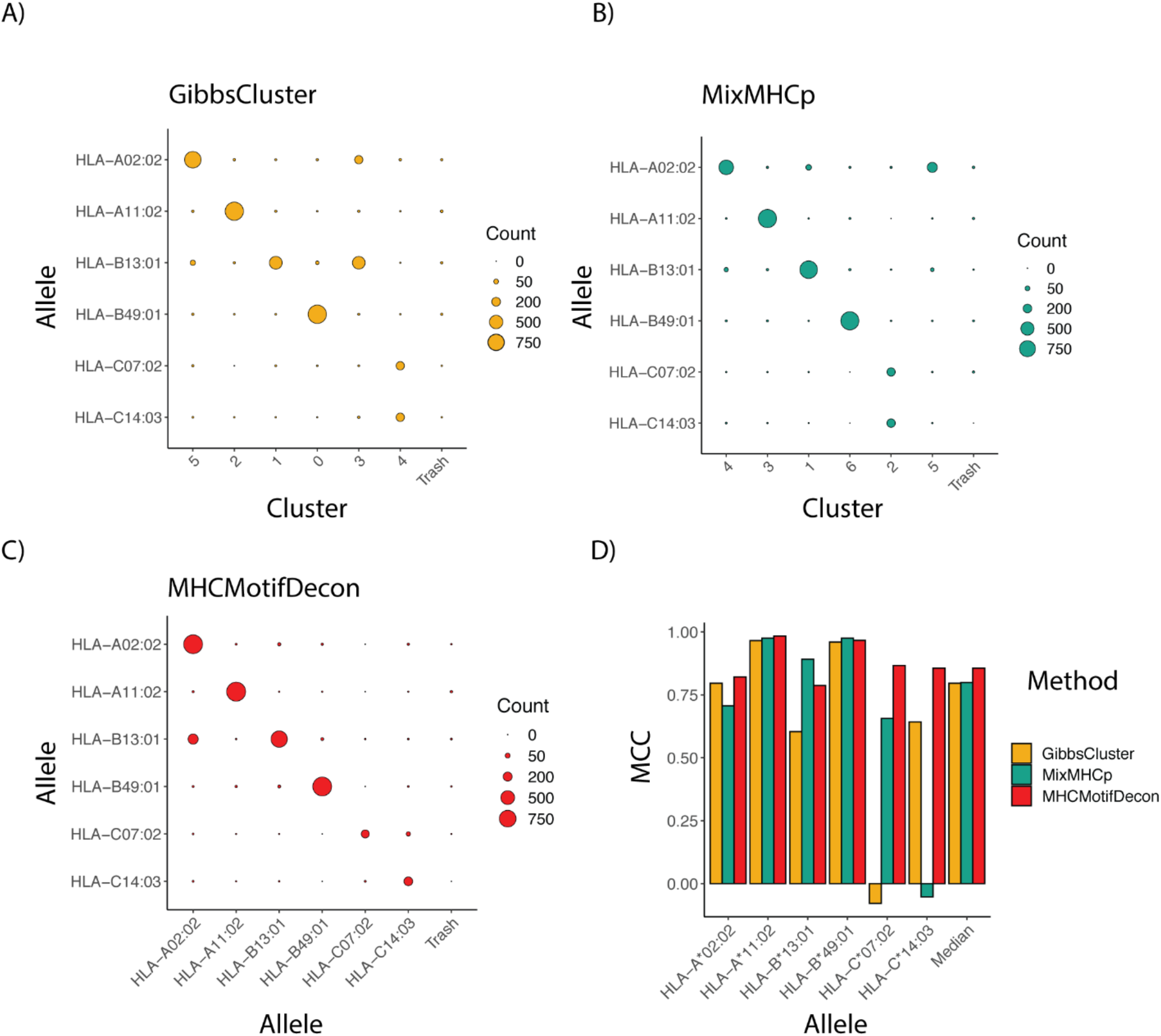
Clustering confusion matrix and Matthews correlation performance for the three methods GibbsCluster, MixMHCp and MHCMotifDecon. Matthews correlation (MCC) performance estimates of accuracy of the motif deconvolution of the different methods for the artificial MS HLA eluted ligand dataset. MCC values for each method and HLA were estimated from the confusion matrices and cluster-HLA annotations shown in figure 1.

To further quantify this, we next estimated, using the HLA association for each cluster as defined in figure 1 and as indicated along the diagonal of each matrix in (figure 2A, B, C), the performance of each method in terms of the Matthews correlation of each HLA/group association. These MCC values for each method and HLA are shown in figure 2D, and confirm the earlier observation with an overall superior performance of MHCMotifDecon and a suboptimal performance of MixMHCp and GibbsClusters for identification of the HLA-C molecule motifs.

### MHC class II deconvolution

To assess the deconvolution efficacy of MHCMotifDecon for HLA class II immunopeptidome data, the method was applied to an artificial dataset generated merging single allele datasets from four different cell lines expressing DRB1*04:03, DRB1*08:03, DRB3*02:02 and DRB5*01:01 alleles respectively (for details on the artificial dataset generation refer to materials and methods section). Here, the motif deconvolution was performed using the new retrained version 4.1 of NetMHCIIpan. The retraining was performed as described in materials and methods expanding the data to include an extended set of single allele MS data, excluding single allele data from the alleles used on the benchmark to avoid overestimation of the deconvolution accuracy. Also in this benchmark two alternative methods, GibbsCluster and MoDec were included. The result of the benchmark is shown in figure 3, again confirming the superior performance of MHCMotifDecon over the other two methods.

**Figure 3.**
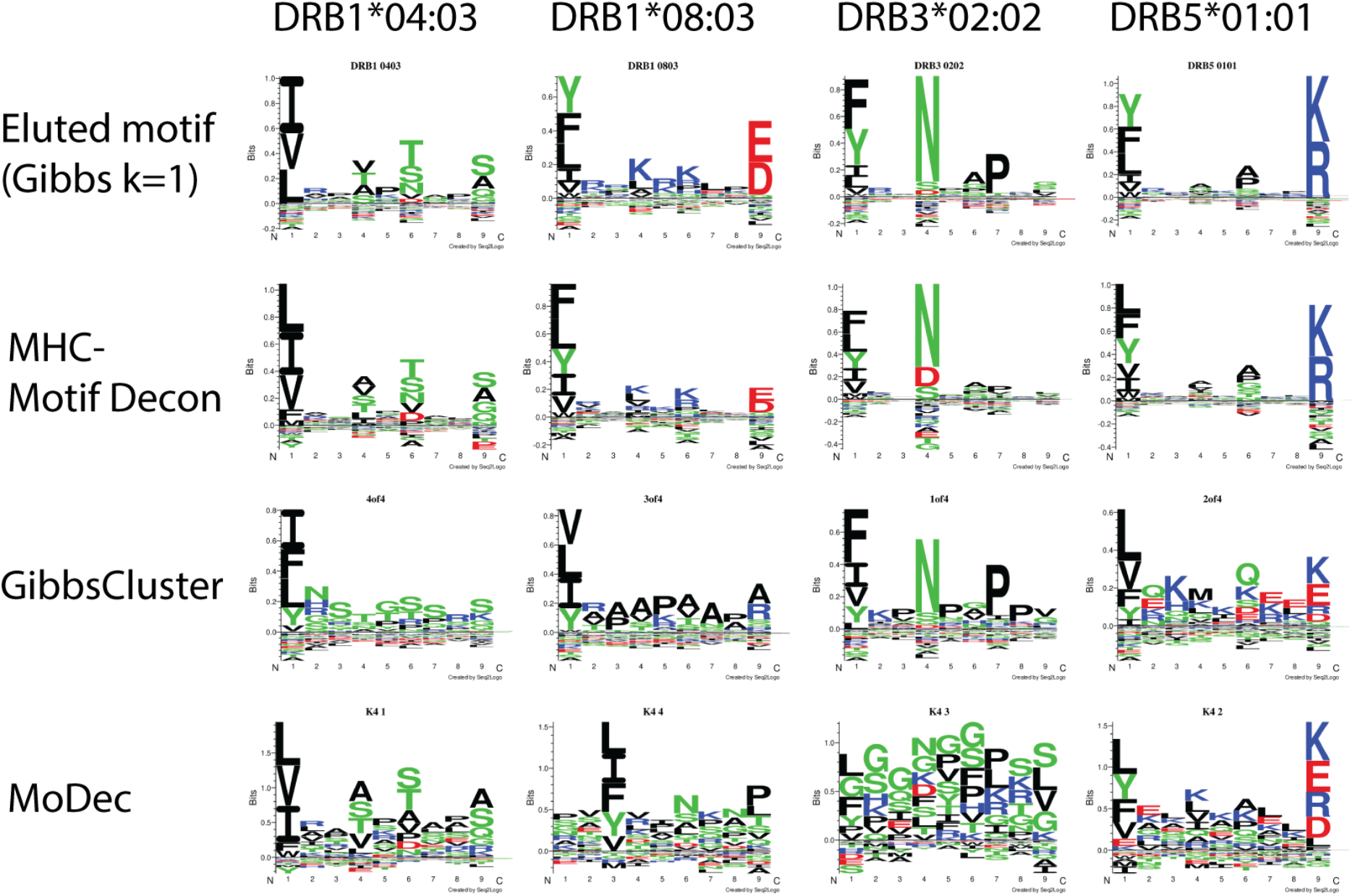
Motif deconvolution of the artificial dataset combining MS eluted ligand data from 4 cell lines each one expressing one individual HLA-DR allele. The three methods MHCMotifDecon, GibbsCluster (Andreatta et al., 2017), and Modec (Racle et al., 2019) were run as described in materials and methods. Eluted motif corresponds to GibbsCluster deconvolution of each of the single allele datasets using a single cluster (k=1).

Similar to class I, a confusion matrix was constructed to quantify the deconvolution power of MHCMotifDecon compared to MoDec and GibbsCluster (figure 4). This figure clearly confirms the superior performance of MHCMotifDecon over the two other methods with a close to 3-fold increase in median performance compared to the second best method (MoDec).

**Figure 4.**
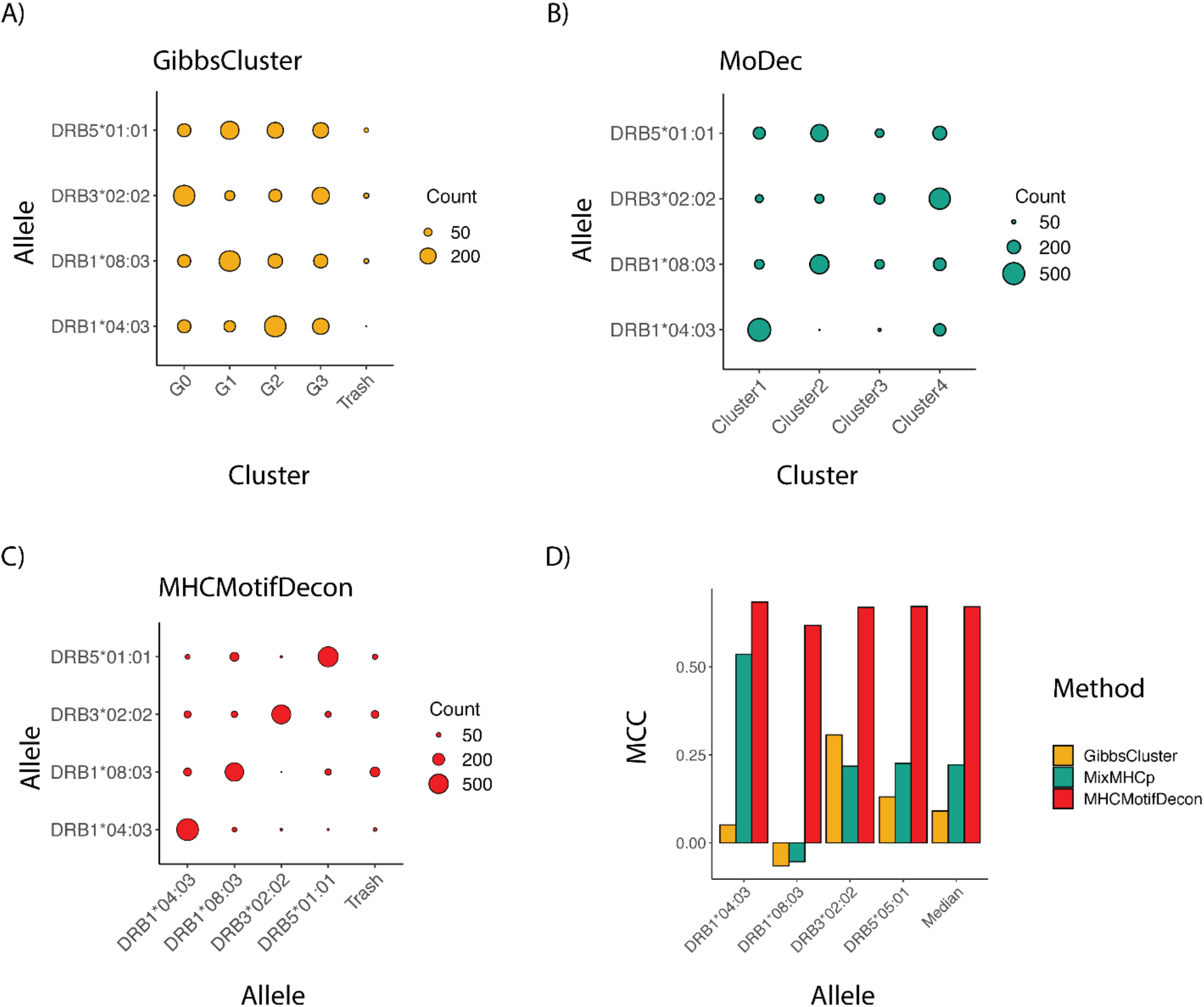
Clustering confusion matrix and Matthews correlation performance for the three methods GibbsCluster, MoDec and MHCMotifDecon. Matthews correlation (MCC) performance estimates of accuracy of the motif deconvolution of the different methods for the artificial MS HLA eluted ligand dataset. MCC values for each method and HLA were estimated from the confusion matrices and cluster-HLA annotations shown in figure 3.

### Dealing with co-immunoprecipitated MS contaminants

MS eluted ligand datasets generally contain a certain amount of co-immunoprecipitated contaminants. One example is the co-immunoprecipitation of varying amounts of HLA class I/II derived peptides that are non-specifically co-purified in addition to the HLA molecule of interest and contaminate the IP preparations (Partridge et al., 2018). This problem that happens despite the use of allele-specific antibodies is inherent to the purification of membrane-bound HLA molecules and is due to the function of the detergent that disrupts the cell membrane and causes cell lysis. As a result, the purified HLA remains bound to a piece of membrane that carries other membrane proteins including other HLA alleles (Partridge et al., 2018). This problem is inevitable unless cells that are engineered to secret the soluble form of the HLA are used (Ben Dror et al., 2010; Trolle et al., 2016) which is not possible in case of tissues and biological samples. To illustrate the power of the tool to handle such contaminants, the complete immunopeptidome data obtained from the IHW09060 (CB6B) cell line expressing the HLA-DR alleles DRB1*13:01 and DRB3*02:02 were submitted to MHCMotifDecon limiting the peptide length to 9-25 and excluding the use of the trash bin option (for details on the data generation refer to materials and methods). The result of this analysis (figure 5A) showed an unusual motif and peptide length distribution for DRB1*13:01 with an additional peak taking up more than half of the peptide counts observed for the shortest peptide length; a clear sign of coimmunoprecipitated contaminants. Rerunning the analysis, setting the trash threshold to 20%, the result in figure 5B was obtained. In this figure, around 33% of the peptides were assigned to the Trash bin, and the motifs and length distribution for the two DR molecules now align with the expectations. To investigate the source of the large proportion of peptides assigned to the Trash bin, the complete IHW09060 peptide data was submitted to the MHCMotifDecon server, now selecting class I, the class I alleles expressed in IHW09060 cell line (HLA-A*01:01,HLA-B*15:01,HLA-C*03:03), and setting a stringent threshold for the trash bin of 2%. The result of the analysis, shown in supplementary figure 2, clearly, and in line with earlier works (Fisch et al., 2021), indicates that the vast majority of the short peptides (in particular for length 8-11) have HLA class I origin. Next, the peptide data was resubmitted for class II motif deconvolution excluding peptides with predicted HLA class I restriction (figure 5C). This analysis confirmed the clear motifs and expected peptide length distributions for the two DR molecules. Also, the analysis confirmed a very minor proportion of remaining MS contaminants (7%), and a trash motif with a highly reduced information content. As a final analysis, the unfiltered data was submitted for class II motif deconvolution limiting the peptide length to 12-25 keeping all other parameters as default. The result of this analysis (figure 5D) aligns closely with the result of the analysis excluding class I restricted peptides (figure 5C) with the exception of the much larger amount of trash peptides found for peptides of length 12 in this latter analysis.

**Figure 5.**
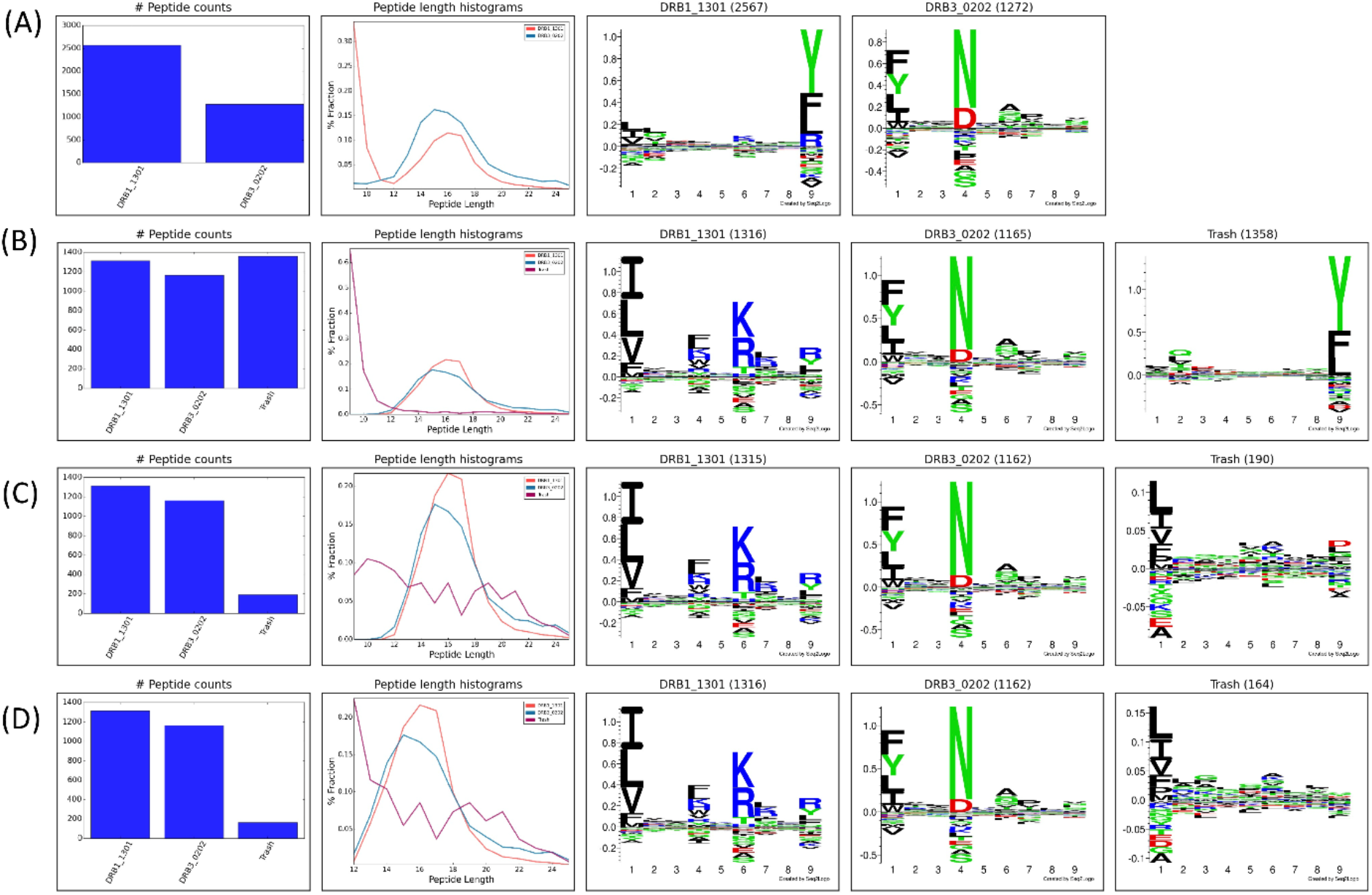
HLA-DR motif deconvolution for the IHW09060 dataset using different strategies to deal with co-immunoprecipitated MS contaminants. A) Peptide length 9-25, excluding the use of the trash bin option (achieved by setting the trash bin threshold to 101%), B) Peptide length 9-25, trash bin 20%, C) Peptide length 9-25, exclude class I binders, trash bin 20%, D) Peptide length 12-25, trash bin 20%.

In summary, this analysis demonstrates the power of MHCMotifDecon to effectively perform HLA motif deconvolution also in immunopeptidome datasets with substantial amounts of copurified contaminants. Furthermore, the analysis suggests that for class II motif deconvolution, placing a peptide length threshold of 12-25 serves as an effective way to deal with class I co-immunoprecipitated contaminants. This however with the caution of potentially identifying a small proportion of false positive identifications at the shortest peptide length. To avoid this, the analysis suggests to filter out potential class I presented peptides prior to performing the class II motif deconvolution.

To further illustrate the issue of potential false positive co-immunoprecipitated MS contaminants, the motif deconvolution of the original data was compared to the deconvolution performed for a set of scrambled HLA-DR ligands (see supplementary figure 3). In the scrambled dataset, the peptide sequences have the same amino acid composition but in a shuffled order (for details refer to methods section). In this comparison, the overlap between rank score in the original and scrambled datasets for HLA class II was found to be substantial for peptide lengths of less than 12 and above 21. In contrast, this overlap is basically absent for peptide length of 12-21 (supplementary figure 3B). Given this, for class II we recommend to filter the peptide length to fall in the range 12-21 when using MHCMotifDecon. Likewise for class I, we recommend applying a length filter to only include peptides of length 8-14 amino acids (see supplementary figure 3A).

### Characterization of the Peptide repertoires of primary and secondary HLA-DR molecules

To better demonstrate the application and power of the MHCMotifDecon method for analysis and interpretation of complex immunopeptidome datasets, a set of 11 HLA-DR immunoprecipitated immunopeptidome datasets were generated from homozygous B cell lines (see supplementary table 1). For details on the data generation refer to materials and methods. Next, this complete set of 11 HLA-DR immunopeptidome was analyzed using MHCMotifDecon with the purpose of characterizing the peptide repertoires of primary and secondary HLA-DR molecules. Note, based on the conclusions from the above analyses, the data were first filtered using MHCMotifDecon to exclude peptides with predicted HLA class I restrictions. Then the peptide data obtained from the 3 cell lines sharing the HLA-DRB3*02:02 allele, were submitted to MHCMotifDecon server 1.0 together with information about their full HLA-DR typing using default MHC class II options and including length histograms and consistency matrix plots. The result of the analysis is shown in figure 6. The result of the analysis of the complete dataset covering all 11 cell line datasets is included in Supplementary figure 4.

**Figure 6.**
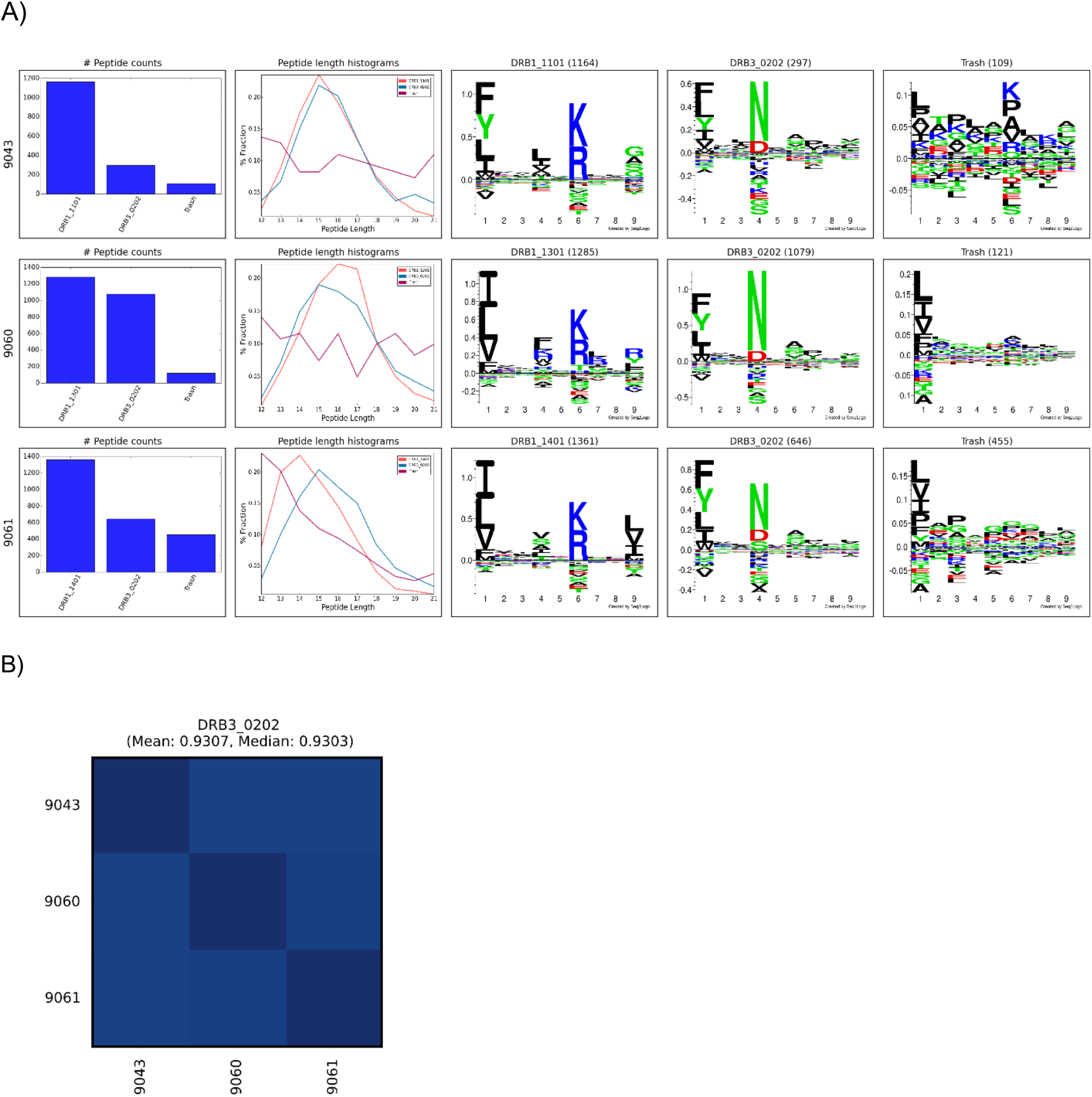
MHCMotifDecon analysis for the peptide data obtained from the three cell lines IHW09043, IHW09060, and IHW09061 all expressing the HLA-DRB3*02:02 allele. A) Peptide counts, length distribution per alleles or trash cluster, and logos from HLA-DR alleles after deconvolution, B) Motif Correlation for HLA-DRB3*02:02 deconvolution in the datasets from the three cell lines IHW09043, IHW09060, and IHW09061. A PCC was calculated from the linearized position specific scoring matrices (PSSM) for the 9 amino acids binding core of each found on the different samples as described earlier (Alvarez et al., 2019).

Overall, these results demonstrated both the high quality of the generated MS HLA-DR immunopeptidome datasets (illustrated by the relative low proportion of peptides assigned to the trash cluster (figure 6A), and a high consistency between the motif deconvolution for HLA-DR molecules shared between multiple cell lines (figure 6B). Also, the figure illustrates the power of the MHCMotifDecon method to accurately deconvolute the DR motifs in these complex datasets (illustrated by the sharp motifs, and the well-defined peptide length distribution for the DR annotated peptides). As a side remark, the analysis further revealed subtle differences in length preference among various primary HLA-DR molecules with DRB1*14:01 showing a preferred length shift towards 14mer peptides, and DRB1*13:01 a shift towards 16mers. These differences will be discussed in more details later.

Based on these encouraging findings, the motif deconvolution of the complete MS HLA-DR immunopeptidome datasets from all 11 cell lines was analyzed in detail in the context of the contribution of the primary and secondary DR molecules to total DR peptidome as well as length distribution of each HLA-DR molecule. In addition, the level of overlap between the DR molecules in each haplotype was determined.

### DRB5 displays a significant contribution to the total immunopeptidome of the HLA-DR51 haplotype group

DRB5 molecules are in linkage disequilibrium with DRB1*15 and DRB1*16 alleles in DR51 haplotype group. The B lymphoblastoid cell lines (SCHU (9013) and CALOGERO (9084)) express the DRB5*01:01 and DRB5*02:02 alleles in combination with DRB1*15 and DRB1*16 and therefore were selected to determine the contribution of DRB5 alleles to HLA-DR51 peptide repertoire. The motif deconvolution of the two cell lines shown in supplementary figure 4 indicates the large contribution of both DRB5 molecules to the peptide repertoire of both cell lines. In fact, DRB5*01:01 and DRB5*02:02 contributed 50% (1490 out of a total of 2991 peptides) and 64% (1010 out of a total of 1580) to the total DR peptidome of 9013 and 9084 respectively (supplementary table 2).

Secondary DR alleles like primary DR molecules show a Bell shape and normal length distribution. When the length distribution of both DRB5 molecules were compared, both showed a preference for the 15 amino acid peptides.

The binding motif of DRB1*15:01 and DRB5*01:01 has been studied before by different groups using crystallography, binding assay and eluted peptides (Li et al., 2000; Reynisson et al., 2020b; Scholz et al., 2017; Smith et al., 1998; Vogt et al., 1994). We were able to refine the motif for the DRB5 alleles of HLA-DR51 haplotype, using a large set of peptides eluted from the homozygous BLCLs and identified by mass spectrometry. In all cases, the identified motif (see supplementary figure 4) was in agreement with previously described motifs (Reynisson et al., 2020b) with some additional residues in the anchor positions P1, P4, P6 and P9.

In DRB5*01:01, **P1** showed preference for bulky aromatic and hydrophobic residues like Tyr (Y) and Trp (W) as well as Phe (F), Leu (L), Ile (I) and Val (V). At **P4** it favors small amino acids like Leu (L), Ala (A), Ile (I) or Val (V). This secondary DR allele accepts amino acids with aliphatic side chain like Ala (A), Ser (S), Gly (G), Pro (P) and Asn (N) at **P6** and positively charged (basic) residues such as Lys (K) or Arg (R) at **P9** that is essential for the interaction between the peptide and the DRB5*01:01 allele.

The same trend was observed in DRB5*02:02. This allele shared the same **P6** and **P9** with DRB5*01:01. However, in DRB5*02:02, **P1** has a strong preference for small amino acids like Ile (I), Val (V) and Leu (L), while **P4** favors the bulky Tyr (Y), Trp (W) and Phe (F) amino acids. Therefore, DRB5*01:01 and 02:02 have different **P1** and **P4** preferences but share the same residues at **P6** and **P9**.

While certain similar features can be identified between the motifs of the primary and secondary DR molecules in the DR51 haplotype, the difference between residue preference at P1 and P4, as well as a distinct basic P9 in both DRB5 molecules, has caused the peptide repertoire of the primary and secondary alleles in the DR51 haplotype to be complementary rather than overlapping (see figure 7 and supplementary table 4).

**Figure 7.**
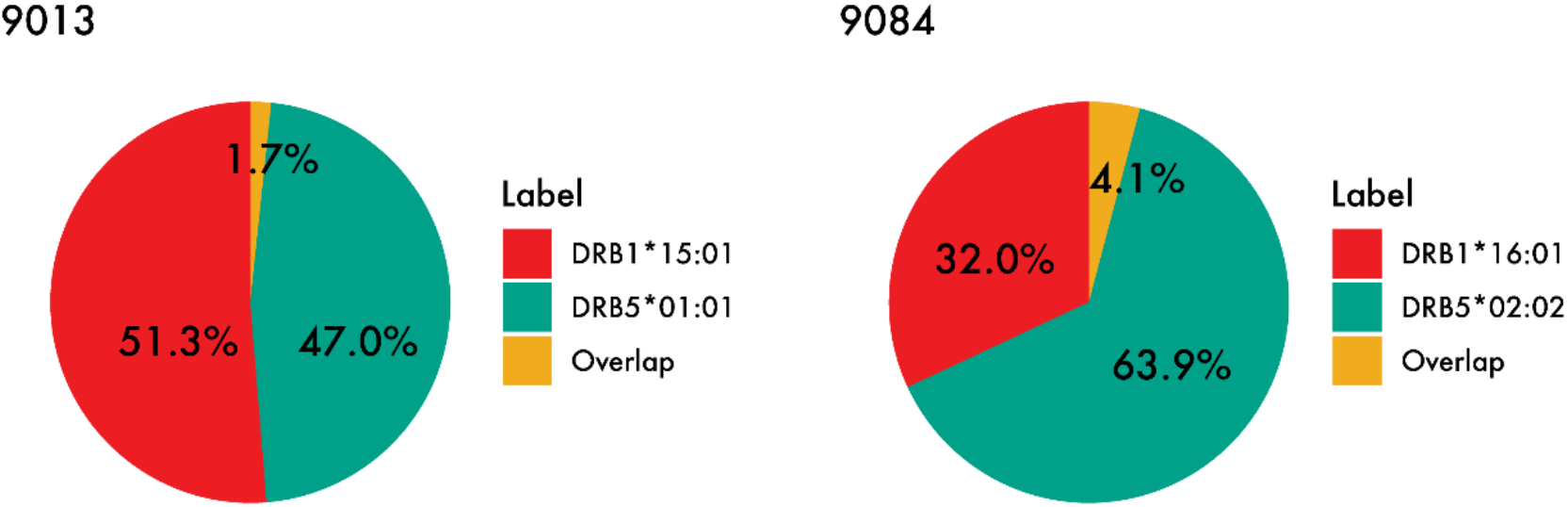
Peptide overlap within the DR51 haplotype group. HLA-DR binding for MS ligands identified within the two DR51 cell lines 9013 (HLA-DRB1*15:01 and HLA-DRB5*01:01) and 9084 (HLA-DRB1*16:01 and HLA-DRB5*02:02) was predicted with NetMHCIIpan-4.1 using a threshold of 1% rank, and the relative contribution from each primary and secondary HLA-DR molecule as well as their overlap were calculated.

### DRB4 has a limited contribution to the DR53 haplotype group peptide repertoire

DRB4 is associated with DRB1*04, DRB1*07 and DRB1*09 in the DR53 haplotype group. Here, we investigated the peptide presentation by these 4 primary and secondary HLA DR using four different cell lines (AWELLS (9090), BER (9093), DKB (9075) and DBB (9052)). The motif deconvolution results in supplementary figure 4 indicate that DRB4 is presenting about 15% of the total DR peptidome. That is, the primary DR molecules (DRB1*04, DRB1*07 and DRB1*09) contribute about 5-6 times more than DRB4 to the total peptide repertoire of DR53 haplotype (see supplementary table 2).

Despite the consistent contribution of DRB4*01:03 in 3 of the cell lines, we observed a below average presentation by this allele in the DBB (9052) cell line (2%, 53 out of a total of 2597 DR annotated peptides). This proportion of presented peptides was comparable to the number of peptides assigned to the trash bin (121), and hence indistinguishable from the noise of the experiment.

Using sequence-based typing method, we noticed a null phenotype for the DRB4*01:03 in this cell line. DRB4*01:03N is characterized by a splice site mutation at the 3’ end of the first intron (AG > AA), leading to a transcript that is larger than that of the common DRB4 molecule and believed to be processed into a non-functional HLA molecule (Voorter et al., 2000). Our data, which is in agreement with the typing results, suggest that this null allele is non-functional or have retained a very limited antigen presentation capacity.

In primary DR molecules, both DRB1*07:01 and DRB1*09:01 showed a preference for shorter peptides (14 mers), while DRB1*04:01 and DRB4*01:03 showed the common class II length preference for 15 amino acid peptides (see supplementary table 3).

The motif for the DRB4*01:03 is distinct as it displays 5 key anchor positions (P1, P4, P6, P7 and P9) instead of the conventional 4 anchor positions (P1, P4, P6 and P9).

**P1** has preference for Leu (L), Ile (I), Val (V), followed by Phe (F) and Met (M). **P4** favors Gln (Q) and Glu (E), and less Val (V), Ala (A) and Leu (L). **P6** accepts amino acids with hydrophobic side chains like Leu (L), Val (V) and Ile (I). **P7** which is unique to DRB4 has strong preference for negatively charged or uncharged polar side chains including Asp (D), Glu (E) and Asn (N) and finally **P9** that shows preference for Gly (G), Ser (S) and Ala (A).

When we compared the motif of the DRB4*01:03 with the associated primary DR alleles (DRB1*04:01, DRB1*07:01 and DRB1*09:01), we noticed that in all combinations the primary DR alleles have a **P1** position that favors the large aromatic residues like Phe (F), Tyr (Y) and Trp (W) (supplementary figure 4). DRB4-restricted epitopes, on the other hand, tend to contain smaller amino acids (Leu, Ile, Val) at the **P1** anchor position.

This lack of similarity in the amino acid preference at P1, which is considered a major determinant of peptide selection, in addition to the 5^th^ anchor position (P7) with strong preference for acidic residues, which is absent in all other DR alleles, explains the difference between the peptide repertoires of primary and secondary DR alleles in DR53 haplotype. In fact, except for DRB1*04:01 and DRB4*01:03 that share some similarity and favor negatively charged residues at P4, the other members of the DRB53 haplotype have little to no similarity in their motifs which results in an extremely low overlap between the ligandome of the DRB1*07 and DRB1*09 with the DRB4*01:03 (see supplementary table 4).

### DRB3 contribution to total DR52 peptide repertoire varies considerably

The DRB3 gene encodes 3 major alleles: DRB3*0101, DRB3*0202, and DRB3*0301 that display a strong association with DRB1*03, DRB1*11, DRB1*12, DRB1*13 and DRB1*14 in the DR52 haplotype family. The cell lines (VAVY (9023), BM21 (9043), CB6B (9060), 31227 ABO (9061) and WT47 (9063)) express frequent variants of DRB3 alleles including DRB3*01:01, DRB3*02:02 and DRB3*03:01 in combination with DRB1*03:01, DRB1*11:01, DRB1*13:01, DRB1*13:02 and DRB1*14:01. We examined the peptide presentation and contribution of different DRB3 molecules to the total DR52 peptide repertoire using above cell lines. We observed a substantial variation in the contribution of the DRB3 molecules to the total DR immunopeptidome ranging from 20 to 56 percent (supplementary table 2). This variation was not only seen among different DRB3 variants but also in one variant (DRB3*02:02) in combination with different primary DR alleles (DRB1*11:01, 13:01 & 14:01). Whether this broad range of contribution is an inherent characteristic of this DRB3 allele or the accompanying primary DR molecules play a role in this, remains to be further investigated.

The overall contribution of DRB3 is more than what was observed for DRB4 (Mean: 15.4%). In some haplotypes, where DRB3 is co-expressed with DRB1*13:01 and DRB1*13:02 alleles, the two DRB3 molecules (DRB3*02:02 and DRB3*03:01) comprise between 46% and 56% of the total DR peptide repertoire respectively, in line with what we observed with DRB5 variants (Mean: 56.8%) (supplementary table 2).

When comparing the length distribution of the 3 DRB3 molecules (supplementary figure 4 and supplementary table 3), we found an interesting pattern in which the DRB3*02:02 showed the normal class II preference for 15 mers whereas a shift toward shorter peptides with a preference for 14 mers was observed for DRB3*01:01 and DRB3*03:01.

In primary DR molecules, we observed a wider range of deviation from the general class II length preference. DRB1*13:01 and DRB1*13:02 showed a preference for longer peptides (16 mers), while DRB1*03:01 and DRB1*11:01 showed the common class II length preference for 15 mers, and DRB1*14:01 favored shorter peptides (14 mers).

Next, we investigated in detail the binding motifs obtained for the three DRB3 molecules included in our dataset. DRB3*01:01 motif consists of only 3 anchor positions (Parry et al., 2007). **P1** in DRB3*01:01 has a preference for large, aromatic amino acids like Tyr (Y) and Trp (W) as well as Phe (F), Leu (L) and Ile (I). At **P4** position, DRB3*0101 shows a distinct preference for Asp (D), which is a negatively charged amino acid. DRB1*0301, which is in linkage disequilibrium with DRB3*01:01 shares the very same preference at **P4**. In crystallography experiments, pocket **P4** has shown a strong positively charged character and therefore the preference for aspartic acid at **P4** is consistent with suggested positive charge of the pocket (Parry et al., 2007).

Due to the extremely small and shallow P6 pocket in the crystal structure, **P6** in DRB3*0101 can only accommodate small residues which has resulted in a “**P1-P4-P9**” binding motif (Parry et al., 2007). **P9** in DRB3*0101, does not show a specific preference and can accept small amino acids like Leu (L), Ile (I), Phe (F) and Val (V).

The DRB3*03:01 molecule shares a series of conserved elements with other DR alleles but has some unique features in the peptide binding groove. In **P1**, it prefers the amino acids with a small aliphatic side chain like Leu (L), Ile (I) and Val (V). This is different from the P1 in the DRB3*01:01, that can accept large aromatic residues like Tyr (Y) and Trp (W) in addition to Phe (F), Leu (L) and Ile (I). At **P4**, DRB3*03:01 has a strong preference for asparagine (N) and aspartic acid (D) which fits in the hydrophilic pocket in the crystal structure of the molecule. In contrast to P4, **P6** does not seem to have preference for any specific amino acid and can accept residues with small side chains like Pro (P), Ala (A), Val (V) and Gly (G). **P9** in both DRB3*01:01 and DRB3*03:01 alleles are very similar and favor small amino acids like Leu (L), Ile (I) and Val (V). P1 and P4 appear to be the key determinants of specific binding in DRB3*03:01 allele (Dai et al., 2008).

The crystal structure of the DRB3*02:02 is not available (Dai et al., 2008). Comparing the motif of this DRB3 variant with the DRB3*01:01 and DRB3*03:01 shows that this molecule shares an identical **P4** with the DRB3*03:01 and displays the same amino acid preference for **P1** as DRB3*01:01. It also shares a similar amino acid composition at **P6** with DRB3*03:01. However, the **P9** in this molecule is different from both DRB3*03:01 and DRB3*01:01 as it favors smaller residues.

Comparison of these three major HLA-DRB3 alleles suggests that they were derived from one another by recombination events that rearranged the four major peptide-binding pockets at peptide positions 1, 4, 6, and 9 (Dai et al., 2008).

When the motif of these DRB3 molecules was compared with their associated primary DR alleles, except for DRB1*03:01 that shares the same preference at **P4** with the DRB3*01:01, the rest of the primary DR molecules showed different preferences in **P1** and **P4** and all presented positively charged residues Lys (K) and Arg (R) at **P6.** Therefore, except for the large overlap that is observed between the peptide repertoire of the DRB1*03:01 and DRB3*01:01, in the rest of the haplotypes the difference between P1 and P4 preferences has caused the peptidome of the DRB3 and the accompanying DRB1 alleles to diverge and the overlap to be minimal (see supplementary table 4).

## Discussion

Analyzing and interpreting complex immunopeptidome data is a highly challenging and intricate task considering the expression of multiple HLA alleles on the cell surface and the presence of co-immunoprecipitated contaminants. Here, we have proposed a simple yet very powerful tool, called MHCMotifDecon, to assist the computationally inexpert users in performing this task. The tool takes the peptide sequences from MS immunopeptidomics experiment(s) together with the associated full HLA typing from one or more samples as input and performs a complete motif deconvolution of the data assigning each peptide to its most likely HLA restriction element while removing probable MS co-immunoprecipitated contaminants.

Using artificial datasets constructed from peptides with known HLA restriction, the method was demonstrated to outperform other available motif deconvolution tools in particular for low expression alleles such as HLA-C in case of HLA class I, and in general across all motifs for HLA class II.

In order to illustrate the application of this tool, the contribution of the primary and secondary DR molecules to total DR peptidome as well as length distribution of each HLA-DR molecule was investigated in detail in a large set of immunopeptidome data generated for this study from a panel of homozygous B lymphoblastoid cell lines (BLCL) selected with an HLA-DR profile that cover the most frequent variants of the secondary DR alleles in the haplotypes with high population coverage. Our results from this analysis show that these secondary DR molecules contribute to the class II DR immunopeptidome far more than what was previously assumed. The ligands presented by secondary DR alleles not only expand the DRB1 peptide repertoire, they might also add an independent function to the haplotype they contribute to (Caillier et al., 2008). Among the secondary DR molecules studied here, DRB5 with the mean contribution of 56.8% demonstrated the highest contribution to the class II ligandome. Next was DRB3 that on average presented (38.5%) of the DR repertoire, followed by DRB4 that displayed the lowest level of peptide presentation (15.4%). Interestingly, we found that in some haplotypes (DR51 and DR52) DRB5 and DRB3 contribute even more than the accompanying DRB1 molecule. DRB4 was the only molecule that consistently contributed less than the DRB1 alleles in the DR53 haplotype family. These findings are in agreement with the previous studies performed on a limited number of haplotypes at transcriptional and mRNA expression level, where, for DRB3 (only DRB3*01:01 and DRB3*02:02 included) the relative proportion of mRNA was reported to vary from being four times lower to almost equal to the amount of accompanying DRB1 mRNA (Faner et al., 2010). DRB5 mRNA was found to be more abundant than the DRB1*150 (Prat et al., 2005), whereas the level of DRB4 mRNA was as much as seven times lower than DRB1*07 (Leën and Gorski, 1997; Stunz et al., 1989).

While there is a considerable degree of variation in the contribution of DRB3 molecules to the DR52 peptidome, these molecules contribute the most when expressed in combination with the DRB1*13:01 and 13:02 primary DRs (46% and 56% respectively). HLA-DRB1*13 alleles have been shown to play a protective role against multiple autoimmune and certain infectious diseases like HBV in HLA-Disease association studies (Bettencourt et al., 2015; Furukawa et al., 2014; Furukawa et al., 2017). It has been hypothesized that the protective role of these DR molecules is due to more efficient antigen presentation and promoting deletion of autoreactive T cells during thymic selection (Bettencourt et al., 2015). However, our results show that the accompanying DRB3 molecule contribute about half of the peptidome when co-expressed with DRB1*13 molecules suggesting that DRB3 molecules that generously add to breadth and diversity of the peptide repertoire, may play a part in the universal protective nature of the DRB1*13 alleles.

On the other end of the spectrum there are DRB1 alleles that are associated with DRB4 molecules. One such allele is DRB1*04:01 that is a risk factor for type 1 diabetes (T1D) and rheumatoid arthritis (RA) (Arango et al., 2017). Our results indicate that DRB4 molecules, due to their limited repertoire, do not contribute to the diversity and extent of the haplotype total peptidome at the same level as DRB3 and DRB5.

Another important finding of this study is that the Secondary DR peptide repertoire complements the peptidome of the DRB1 molecules and is not redundant. The complementarity or redundancy of the peptide repertoires of the DRB3, 4 and 5 alleles has long been a source of controversy and overlap between the peptide repertoires of the 2 DR molecules in a given haplotype has been reported by different groups (Texier et al., 2001; Wucherpfennig et al., 1994). However, our results demonstrate that the overlap between the DR molecules in all the haplotypes studied here is minimal with the only exception being the DRB1*03:01 and DRB3*01:01. These findings are supported by differences in the motifs and amino acid preferences in different anchor positions. Some of these differences can be seen consistently in all haplotypes and are not allele specific like P1 and P4 pockets that exhibit distinct binding preferences.

The P1 pocket in the crystal structure of the HLA-DR molecules is dimorphic and contains either glycine or valine at position 86 of the B chain. Presence of glycine produces a larger pocket which results in a preference for bulky hydrophobic and aromatic residues. Substitution of glycine with valine creates a shallow hydrophobic pocket that is only able to accommodate small hydrophobic amino acids (Gupta et al., 2008; Hammer et al., 1995). Intriguingly we have found that in all haplotypes studied here, except for (DRB1*11:01, DRB3*02:02), where one DR allele (either primary or secondary) has a large P1 pocket and prefers the bulky aromatic residues like Tyr (Y) and Trp (W), the second DR allele favors the small aliphatic residues like Ile (I), Lue (L) and Val (V). This is in agreement with P1 pocket role in determining allele specific preferences for anchor residues (Gupta et al., 2008) and explains how this structural and steric hindrance at P1 pocket causes the primary and secondary DR alleles to present diverse peptidome with minimal overlap.

The P4 pocket is the most diverse and polymorphic pocket of the HLA-DR binding site (Smith et al., 1998) which results in the different residue preferences at P4 in the peptide repertoire of DR molecules in a haplotype. This is another factor that contributes to the presentation of distinct peptidomes by primary and secondary DR molecules. The significance and impact of the position 4 in creating a diverse peptidome is even more highlighted when we appreciate the large degree of overlap between DRB3*01:01, DRB1*03:01 which is mainly due to sharing the identical P4 anchor residue Asp (D) for the two molecules. It has been suggested that the P4 pocket of HLA-DR makes an important contribution to susceptibility to different human autoimmune diseases (Smith et al., 1998).

In addition to the distinct residue preferences at P1 and P4 that cause the peptidome of the primary and secondary DR alleles to be different from each other in all haplotypes, there are other distinguishing features that are unique to certain haplotypes. For example, in the DR53 haplotype, the additional anchor position (P7) in DRB4*01:03, imposes very different peptide binding requirements between the primary and secondary DR molecules explaining why DRB4 displays the least number of overlapping peptides with the accompanying DR7 and DR9 molecules.

In the end, the non-redundant and complementary nature of the secondary DR peptidomes is not unexpected as throughout the evolution different types of HLA have been developed by recombination events, to add diversity to the immune system and better defend the body against external pathogens. Our data indicate that the secondary DR peptide repertoire in all haplotypes largely expands the peptidome of the DRB1 molecules (James et al., 2018; Scholz et al., 2017). More importantly these peptides similar to their accompanying DRB1 presented peptides can activate autoreactive CD4^+^ T cells and provoke an autoimmune response which has been demonstrated by multiple studies where the DRB3, 4 and 5 restricted T cells are detected in measurable amount in patients with infectious and autoimmune diseases (James et al., 2018; Wang et al., 2020).

In summary, given the extent of contribution of the secondary DR alleles, their complementary and non-redundant nature, in addition to their ability to successfully provoke a T cell response reveal the significance of these molecules and strongly suggest that they should be regarded as functionally independent alleles in future studies of autoimmune and infectious diseases.

The proposed MHCMotifDecon tool is supervised and performs the motif deconvolution by use of predicted HLA restrictions. While this, as demonstrated here, provides a highly powerful and successful approach for motif deconvolution and handling of complex MS peptidomics datasets, one should bear in mind that these successes are contingent on accurate binding prediction power being available for the MHC molecules in question. Also, one should note that the tool makes unambiguous assignment of HLA restrictions, hence failing to report multi-allele binding of promiscuous peptides.

With these cautions in mind, in conclusion, we have proposed a novel and powerful tool allowing computationally novice users to analyze and interpret complex immunopeptidome datasets, enabling accuracy assessment, motif deconvolution and HLA restriction assignment, estimation of relative allele contribution, and length profile investigations. The tool was demonstrated to achieve state-of-the-art performance and illustrated to enable novel biological discoveries. Given its ease of use, we expect the tool to be highly valuable for the scientific community facilitating better use and understanding of immunopeptidome data.

## Acknowledgements

We would like to thank Dr. Rico Buchli (Pure Protein, LLC) for providing the L243 affinity columns for this study. We sincerely thank Steven Cate (University of Oklahoma Health Sciences Center) and Sean Osborn (Pure MHC, LLC) for HLA typing of the BLCLs and very helpful discussions. This research was funded in part through US Federal funds from the National Institute of Allergy and Infectious Diseases, National Institutes of Health and Department of Health and Human Services (under Contract No. HHSN272201200010CERC to M.N).

## Author Contributions

Conceptualization, S.K., C.B., W.H.H, and M.N.; Methodology, S.K., C.B., B. A., and M.N; Investigation, S.K., C.B., B. A., H.Y., and M.N.; Writing – Original Draft, S.K., C.B., H.Y., W.H.H, and M.N.; Writing – Review & Editing, S.K., C.B., and M.N.; Funding Acquisition and Resources, W.H.H, and M.N.; Supervision, S.K., C.B, and M.N.

## Materials and Methods

### MHCMotifDecon method

Input peptides are first filtered by length range (determined by the user) with default lengths for MHC class I, 8-14, and for MHC class II, 13-21. All the remaining unique peptides are then predicted for MHC presentation towards all the MHC alleles expressed in the given cell line using NetMHCpan4.1 (Reynisson et al., 2020a) or NetMHCIIpan (a retrained version for this paper). NetMHCpan and NetMHCIIpan methods can make a prediction for all MHC molecules with known protein sequence. A key prediction value from the two methods is the percentile rank score for the likelihood of a peptide being presented by a given MHC molecule. This score reflects the likelihood of a random natural peptide obtaining a prediction score equal to or higher than the score for the given peptide. This score hence ranges from 0-100%, where 0 corresponds to the strongest possible score exceeding the score for all random natural peptides. For MHC class I, a percentile rank value of 2% or lower is considered a binder, and for class II the corresponding value is 5% (Reynisson et al., 2020a). Likewise, earlier work has shown that peptides with a percentile rank score > 20% are MS co-immunoprecipitated contaminants (Reynisson et al., 2020b). Given this, by default all the peptides with a higher rank score of 20% to all the proposed alleles are considered as a non-binder or contaminant, and therefore assigned to the trash cluster. Optionally, threshold can also be tuned by the user to be more or less stringent.

The graphical output of the tool provides: sequence motif logos for all the present alleles with a minimum of 10 (default value) peptides assigned to it, the length distribution for all the alleles and the trash cluster, and a bar-plot with peptide counts assigned to each of the clusters. In addition, a correlation matrix can be plotted when analyzing several samples with the same HLA alleles on the same run, to check for consistency.-The correlation matrices are then calculated by stacking all the amino acid frequencies for all the core positions in a vector of 20*9 and obtaining a Pearson correlation coefficient (PCC) among the method information and the sample data. Additional text datafiles provided in the output can be downloaded after the analysis and include the raw and rank scores for each peptide-allele combination, and the 9-mer core peptide sequence. We provide this tool as a web server at https://services.healthtech.dtu.dk/service.php?MHCMotifDecon-1.0.

### Retrain of NetMHCIIpan

NetMHCIIpan was specifically retrained for this publication (updated version called NetMHCIIpan-4.1) excluding all single allele data from DRB1*04:03, DRB1*08:03, DRB3*02:02 and DRB5*01:02 alleles. Single allele data for those alleles were left out to be used for the independent benchmark of MHCMotifDecon. Compared to NetMHCIIpan-4.0, additional single allele datasets from Abelin et al. publication were included (Abelin et al., 2019), and the binding affinity data were re-curated to include updated data for DQ4. The data used to retrain this method is available at http://www.cbs.dtu.dk/suppl/immunology/NetMHCIIpan-4.1. Apart from the modified data, the method is identical to the previously published version of NetMHCIIpan 4.0 (Reynisson et al., 2020a). A performance comparison of NetMHCIIpan-4.1 and 4.0 is included in supplementary figure 1 demonstrating a significantly improved performance of version 4.1. NetMHCIIpan-4.1 is available at https://services.healthtech.dtu.dk/service.php?NetMHCIIpan-4.1.

### Benchmark Datasets

#### Artificial MS MHC class I eluted ligand dataset

Peptides of length 8-14 amino acids were collected from (Abelin et al., 2017). Peptides sharing an 8mer or longer overlap with the NetMHCpan-4.1 training data were excluded. A random subset of 1000 peptides were sampled from the HLA-A*02:02, HLA-A*11:02, HLA-B*13:02 and HLA-B*49:01 single allele datasets, and 200 peptides were sampled from HLA-C*07:02, and HLA-C*14:03 datasets.

#### Artificial MS MHC class II eluted ligand dataset

Peptides with single allele associations for MHC class II were retrieved from MassIVE, MSV000083991 (Abelin et al., 2019). Eluted ligands from cell lines expressing only one of the alleles DRB1*04:03, DRB1*08:03, DRB3*02:02 and DRB5*01:02 were used to generate the artificial dataset for class II. The four single allele datasets were excluded from NetMHCIIpan retraining performed for this paper, to avoid overlapping peptides present on the test and the train set. Additionally, peptides that shared a stretch of 9 amino acids with the dataset used for training NetMHCIIpan were excluded. Finally, for each of the alleles mentioned before, 800 unique peptides of length 13-21 were combined to generate the artificial dataset for MHC-II.

### Deconvolution of immunopeptidome data

MHCMotifDecon was used with the default parameters for class I artificial dataset (Length range = 8-14; Class = I; Threshold for trash cluster = 20) and for class II artificial dataset (Length range = 13-21; Class = II; Threshold for trash cluster = 20). For the in-house MHC class II datasets, the length range was expanded to 12-21 to further analyze length specificities.

GibbsCluster deconvolution for MHC class I artificial dataset was performed using default parameters as described earlier (Andreatta et al., 2017), and including 1-6 clusters, motif length 9 amino acids, using “trash cluster” with a threshold=2, performing single sequence moves at every iteration, max length deletions = 5, max length insertions=1, number of seeds for initial starting conditions=5, Number of iterations x sequence x temperature step = 50.

The deconvolution softwares MixMHCp and MoDec were downloaded from the GitHub repositories (https://github.com/GfellerLab/MixMHCp and https://github.com/GfellerLab/MoDec) and run with default parameters for class I and class II (Bassani-Sternberg and Gfeller, 2016; Racle et al., 2019).

As MHCMotifDecon by construction will identify clusters equal to the HLA alleles defined in the sample, the number of clusters for the deconvolution of the artificial immunopeptidome datasets was set to six for MHC class I and four for MHC class II for MixMHCp and GibbsCluster methods included in the comparison. Also, as MixMHCp and GibbsCluster only perform motif clustering and do not assign HLA allele to the identified clusters, the HLA assignment for these methods was performed by visual inspection in unambiguous cases and by performance optimization in ambiguous cases.

### Scrambled dataset generation

All MS peptide data from eleven cell lines in the in-house MHC class II datasets were merged (N=49,550). Each peptide sequence was scrambled 100 times and predicted with NetMHCIIpan-4.1 for all the alleles expressed by the cell line presenting the original peptide sequence. For each peptide-scramble combination the best (lowest) rank score was selected. A random sample of the scrambled peptide data was used to match the counts of the original dataset following the same length distribution.

### Performance metrics and statistical significance

A Matthews coefficient correlation (MCC) was calculated for each prediction method and allele, from the confusion matrix constructed from the complete motif deconvolution matrix (like the ones shown in figure 3) assigning the true positive count (TP) as the number of peptide sequences from the Allele_X dataset assigned to Cluster_X, the false positives count (FP) to the number of peptides assigned to Cluster_X but belonging to any of the remaining alleles in the sample and excluding the trash cluster. Finally, the false negative (FN) count was assigned as the number of peptides from the Allele_X dataset assigned to any of the remaining Clusters, and the true negative (TN) count as the number of peptides that were correctly predicted as non-binders to Allele_X. For GibbsCluster, MixMHCp and MoDec that do not provide an allele association, the cluster with the majority of peptides assigned to that allele was used. In cases where the majority of peptides on that cluster were assigned to a previously assigned allele (example Cluster 3 in figure 3A) the cluster was assigned to the remaining and non-assigned allele (DRB5*01:01 allele in the previous example). After all allele-specific MCC were calculated, the median was used to compare MCCs across methods.

Bootstrapping samples of N=100 with repetitions were used to add statistical significance to the random associations.

### Sequence Logo motifs

All the logos in this publication were constructed from the 9-mer cores provided by each deconvolution algorithm and using Seq2Logo without clustering and using P-weighted Kulblack-Leibler option (Thomsen and Nielsen, 2012). Additionally, MHCMotifDecon web server provides sequence logos of the deconvoluted motifs using the same software with default options.

### Cell lines and antibody

Homozygous B lymphoblastoid cell lines (BLCL) were obtained from the International Histocompatibility Working Group (IHWG) Cell and DNA bank housed at the Fred Hutchinson Cancer Research Center, Seattle, WA (http://www.ihwg.org). A group of 11 cell lines expressing the high frequency DRB3, DRB4 and DRB5 molecules along with the DRB1 alleles were selected for the study (supplementary table 1). To guarantee intact Class II processing and presentation machinery and to ensure that the total HLA-DR expression and the ratio of the primary and secondary DR alleles represent the physiological level use of engineered cells was avoided.

The cells were grown in high density cultures in roller bottles in complete RPMI medium (Gibco) supplemented with 15% fetal bovine serum (FBS; Gibco/Invitrogen Corp) and 1% 100 mM sodium pyruvate (Gibco). Cells maintained >90% viability at all times during the culture and were harvested from the suspension when the cell density reached to 1.5 −2.5e6 cells/ml. The cells were washed twice with ice cold PBS and spun down at 2500xg at 4C for 10 minutes. The cell pellets were snap frozen in LN2 and stored at −80 until downstream processing.

All cell lines were subjected to high-resolution sequence-based HLA typing (HLA-A, -B, -C, DRB1,3, 4, 5, DP and DQ) immediately upon receipt and growth in our laboratory, for authentication prior to large scale culture and data collection.

### Antibody

The anti-human HLA-DR antibody (clone L243) was used for extraction of total HLA DR from the BLCLs. The L243 monoclonal antibody reacts with the HLA-DR antigen and does not cross react with HLA-DP and HLA-DQ. Clone L243 binds a conformational epitope on HLA-DRα which depends on the correct folding of the αβ heterodimer and therefore can be used to purify different DR molecules regardless of the DRB chain.

### Isolation and purification of HLA-DR bound peptides

HLA-DR molecules were purified from homozygous cell lines by affinity chromatography using the mAb L243 (American Type Culture Collection, Manassas, VA) coupled to CNBr-activated Sepharose 4 Fast Flow (Amersham Pharmacia Biotech, Orsay, France) as described previously with some modifications (Purcell et al., 2019). Briefly, frozen cell pellets were pulverized using Retsch Mixer Mill MM400, resuspended in lysis buffer comprised of Tris pH 8.0 (50 mM), octylphenoxy poly (ethyleneoxy) ethanol (Igepal, 0.5%), NaCl (150 mM) and complete protease inhibitor cocktail (Roche, Mannheim, Germany). Lysates were centrifuged at 200,000 xg for 90 min in an Optima XPN-80 ultracentrifuge (Beckman Coulter, IN, USA) and filtered supernatants were loaded on immunoaffinity columns. After a minimum of 3 passages, columns were washed sequentially with a series of wash buffers (Purcell et al., 2019) and were eluted with 0.2 N acetic acid. The HLA was denatured, and the peptides were isolated by adding glacial acetic acid (up to 10%) and heat. The mixture of Peptides and HLA-DR was subjected to reverse phase high performance liquid chromatography (RP-HPLC).

### First-dimension liquid Chromatography

Reverse-phase high performance liquid chromatography (RP-HPLC) was used to reduce the complexity of the peptide mixture eluted from the affinity column. First, the eluate was dried under vacuum by using a CentriVap concentrator (Labconco, Kansas City, Missouri, USA). The solid residue was dissolved in 10% acetic acid in water and fractionated over a 150-mm long Gemini C_18_ column, pore size 110 Å, particle size 5μ (Phenomenex, Torrance, California, USA) by using a Paradigm MG4 instrument (Michrom BioResources, Auburn, California, USA). An acetonitrile (ACN) gradient was run at pH 2 using a two-solvent system. Solvent A contained 2% ACN in water, and solvent B contained 5% water in ACN. Both solvent A and Solvent B contained 0.1% trifluoroacetic acid (TFA). The column was pre-equilibrated at 2% solvent B. After loading the peptide mixture on the column in a period of 19 min by using a solvent system comprised of 2% Solvent B with the flow rate of 160, two linear gradients were run at the same flow rate: 4% to 40% Solvent B for 40 min, followed by 40% to 80% Solvent B for 8 min. The percentage of Solvent B was maintained at 80% for 4 min, and then decreased to 2% over a period of 3 min. Fractions were collected in 2 min intervals using a Gilson FC 203B fraction collector (Gilson, Middleton, Wisconsin, USA), and the ultra-violet (UV) absorption profile of the eluate was recorded at 215 nm wavelength.

### Second-dimension liquid chromatography

Peptide-containing first dimension HPLC fractions were dried, resuspended in an aqueous solvent composed of 10% acetic acid, 2% ACN and iRT peptides (Biognosys, Schlieren, Switzerland) as internal standards. Fractions were applied individually to an Eksigent nanoLC 415 nanoscale RP-HPLC (AB Sciex, Framingham, Massachusetts, USA), including a 0.5-mm long, 350 mm internal diameter Chrom XP C18 trap column with 3-mm particles and 120Å pores, and a 15-cm-long ChromXP C18 separation column (75-mm internal diameter) packed with the same medium (AB Sciex, Framingham, Massachusetts, USA). An ACN gradient was run at pH 2.5 using a two-solvent system. Solvent A was 0.1% formic acid in water, and solvent B was 0.1% formic acid in 95% ACN in water. The column was pre-equilibrated at 2% solvent B. Samples were loaded at 5 μL/min flow rate onto the trap column and run through the separation column at 300 nL/min with two linear gradients: 10% to 40% B for 70 minutes, followed by 40% to 80% B for 7 minutes.

### MS

The column effluent was injected using the nanospray III ion source of an AB Sciex TripleTOF 5600 quadrupole time-of-flight mass spectrometer (AB Sciex, Framingham, MA, USA) with the source voltage set to 2,400 v. Information-dependent analysis (IDA) of peptide ions was acquired based on a survey scan in the TOF-MS positive-ion mode over a range of 300 to 1,250 m/z for 0.25 seconds. Following each survey scan, up to 22 ions with a charge state of 2 to 5 and intensity of at least 200 counts per second were subjected to collision-induced dissociation (CID) for tandem MS analysis (MS/MS) over a maximum period of 3.3 seconds. Selection of a particular ion m/z was excluded for 30 seconds after three initial MS/MS experiments. Dynamic collision energy was utilized to automatically adjust the collision voltage based upon ion size and charge. PeakView Software version 1.2.0.3 (AB Sciex, Framingham, MA, USA) was used for data visualization.

### Data analysis

Peptide sequences were identified using PEAKS Studio 10.5 software (Bioinformatics Solutions, Waterloo, Canada) at a precursor mass error tolerance of 30 ppm and a fragment mass error tolerance of 0.02 Da. A database composed of SwissProt *Homo sapiens* (taxon identifier 9606) and iRT peptide sequences was used as the reference for database search. Variable post-translational modifications (PTM) including acetylation, deamination, pyroglutamate formation, oxidation, sodium adducts, phosphorylation, and cysteinylation were included in database search. Identified peptides were further filtered at a false discovery rate (FDR) of 1% using PEAKS decoy-fusion algorithm.

### Peptide Repertoire Overlap Analysis

To determine the degree of overlap between the peptide repertoire of the primary and secondary DR molecule in each haplotype, the total DR peptide repertoire of each cell line was submitted to NetMHCIIpan 4.1 and the predicted binding to the primary and secondary molecule was calculated. Peptide binders were defined using an EL rank threshold of 1%, peptides that showed binding to both molecules were considered overlapping. Note, that this analysis was performed using a different predicted binding threshold (1%) compared to that used by MHCMotifDecon (20%). This is because the latter analysis was performed to exclude the co-purified contaminants, while the former was done to compare the overlap between highly reliable peptide repertoires. Therefore, the reported relative repertoire sizes of the primary and secondary DR molecule can be slightly different between the two analyses.

