## Supplementary information for "Accurate MHC Motif Deconvolution of Immunopeptidomics Data Reveals a Significant Contribution of DRB3, 4 and 5 to the Total DR Immunopeptidome"

**Supplementary Table 1**

| Haplotype Group | IHW ID | Alternate ID | Class I HLA | | | Class II HLA (DR) | | | |
|---|---|---|---|---|---|---|---|---|---|
| | | | A | B | C | DRB1 | DRB3 | DRB4 | DRB5 |
| **DR52** | **IHW09023** | VAVY | 01:01 | 08:01 | 07:01 | 03:01 | 01:01 | | |
| | **IHW09043** | BM21 | 01:01 | 41:01 | 17:01 | 11:01 | 02:02 | | |
| | **IHW09060** | CB6B | 01:01 | 15:01 | 03:03 | 13:01 | 02:02 | | |
| | **IHW09061** | 31227ABO | 02:01 | 18:01 | 07:01 | 14:01* | 02:02 | | |
| | **IHW09063** | WT47 | 32:01 | 44:02 | 05:01 | 13:02 | 03:01 | | |
| **DR53** | **IHW09052** | DBB | 02:01 | 57:01 | 06:02 | 07:01 | | 01:03N | |
| | **IHW09075** | DKB | 24:02 | 40:01 | 03:04 | 09:01 | | 01:03 | |
| | **IHW09090** | AWELLS | 02:01 | 44:02 | 05:01 | 04:01 | | 01:03 | |
| | **IHW09093** | BER | 02:01 | 13:02 | 06:02 | 07:01 | | 01:03 | |
| **DR51** | **IHW09013** | SCHU | 03:01 | 07:02 | 07:02 | 15:01 | | | 01:01 |
| | **IHW09084** | CALOGERO | 02:01 | 40:02 | 02:02 | 16:01 | | | 02:02 |

HLA type of the Homozygous BLCLs including Class I (A, B, C) and Class II (DRB1, 3, 4 5)

*:  Not sequenced with exon3 for possible DRB1*14:54

**Supplementary Table 2**

| Haplotype Group | Cell Line IHW ID | Primary DR | Count | Proportion | Secondary DR | Count | Proportion |
|---|---|---|---|---|---|---|---|
| **DR52** | 9023 | DRB1*03:01 | 1545 | 61.6% | DRB3*01:01 | 964 | 38.4% |
| | 9043 | DRB1*11:01 | 1164 | 79.7% | DRB3*02:02 | 297 | 20.3% |
| | 9060 | DRB1*13:01 | 1285 | 54.4% | DRB3*02:02 | 1079 | 45.6% |
| | 9061 | DRB1*14:01 | 1361 | 67.8% | DRB3*02:02 | 646 | 32.2% |
| | 9063 | DRB1*13:02 | 964 | 43.8% | DRB3*03:01 | 1237 | 56.2% |
| **DR53** | 9052 | DRB1*07:01 | 2597 | 98% | DRB4*01:03N | 53 | 2% |
| | 9075 | DRB1*09:01 | 1575 | 85.4% | DRB4*01:03 | 270 | 14.6% |
| | 9090 | DRB1*04:01 | 2360 | 83.9% | DRB4*01:03 | 453 | 16.1% |
| | 9093 | DRB1*07:01 | 1751 | 84.4% | DRB4*01:03 | 323 | 15.6% |
| **DR51** | 9013 | DRB1*15:01 | 1501 | 50.2% | DRB5*01:01 | 1490 | 49.8% |
| | 9084 | DRB1*16:01 | 570 | 36.1% | DRB5*02:02 | 1010 | 63.9% |

Peptide contribution of the primary and secondary HLA-DR molecules as obtained by MHCMotifDecon.

**Supplementary Table 3**

| Haplotype Group | Cell Line IHW ID | Primary DR | Mode | Secondary DR | Mode |
|---|---|---|---|---|---|
| **DR52** | 9023 | DRB1*03:01 | 15 | DRB3*01:01 | 14 |
| | 9043 | DRB1*11:01 | 15 | DRB3*02:02 | 15 |
| | 9060 | DRB1*13:01 | 16 | DRB3*02:02 | 15 |
| | 9061 | DRB1*14:01 | 14 | DRB3*02:02 | 15 |
| | 9063 | DRB1*13:02 | 16 | DRB3*03:01 | 14 |
| **DR53** | 9052 | DRB1*07:01 | 14 | DRB4*0103N | 15 |
| | 9075 | DRB1*09:01 | 14 | DRB4*01:03 | 15 |
| | 9090 | DRB1*04:01 | 15 | DRB4*01:03 | 15 |
| | 9093 | DRB1*07:01 | 14 | DRB4*01:03 | 15 |
| **DR51** | 9013 | DRB1*15:01 | 15 | DRB5*01:01 | 15 |
| | 9084 | DRB1*16:01 | 16 | DRB5*02:02 | 15 |

Prevalent length (mode) for the peptides mapped to the primary and secondary HLA-DR molecules. Mode values are obtained from the peptide length distributions included in supplementary figure 4.

**Supplementary Table 4**

| Haplotype Group | Cell Line IHW ID | Overlap | Primary DR | Proportion | Secondary DR | Proportion |
|---|---|---|---|---|---|---|
| DR52 | 9023 | 27.5% | DRB1*03:01 | 44.8% | DRB3*01:01 | 27.7% |
| | 9043 | 1.5% | DRB1*11:01 | 73.1% | DRB3*02:02 | 25.4% |
| | 9060 | 0.9% | DRB1*13:01 | 37.0% | DRB3*02:02 | 62.1% |
| | 9061 | 3.5% | DRB1*14:01 | 59.3% | DRB3*02:02 | 37.2% |
| | 9063 | 11.0% | DRB1*13:02 | 29.4% | DRB3*03:01 | 59.6% |
| DR53 | 9052 | 0.8% | DRB1*07:01 | 98.4% | DRB4*01:03N | 0.8% |
| | 9075 | 0.6% | DRB1*09:01 | 80.3% | DRB4*01:03 | 19.2% |
| | 9090 | 7.0% | DRB1*04:01 | 75.8% | DRB4*01:03 | 17.2% |
| | 9093 | 0.3% | DRB1*07:01 | 78.5% | DRB4*01:03 | 21.2% |
| DR51 | 9013 | 1.7% | DRB1*15:01 | 51.3% | DRB5*01:01 | 47.0% |
| | 9084 | 4.1% | DRB1*16:01 | 32.0% | DRB5*02:02 | 63.9% |

**The proportion of the DR ligandome predicted to be presented by either the primary (DRB1), secondary (DRB3, 4, or 5) or by both DR molecules.** HLA presentation was predicted using NetMHCIIpan-4.1 with a rank threshold of 1%.

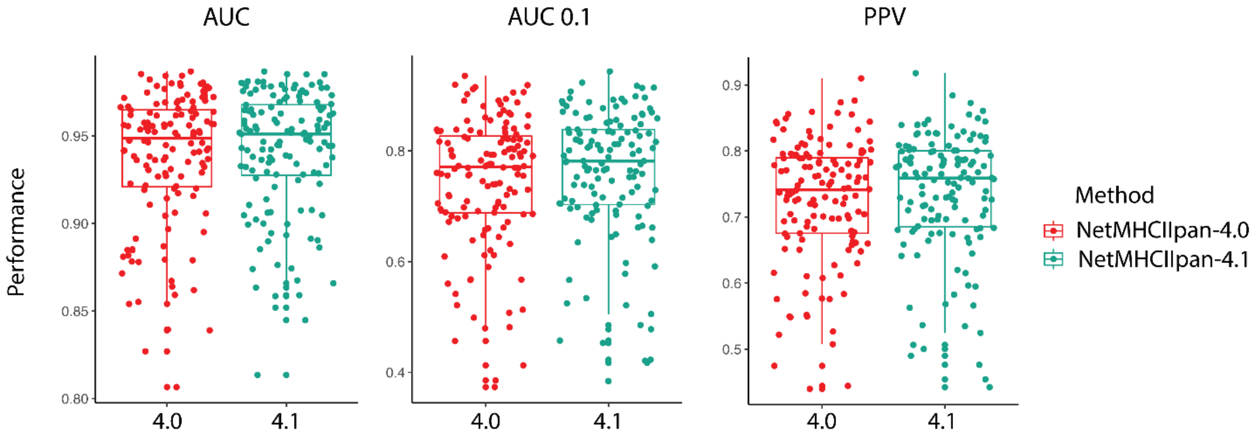


**Supplementary Figure 1.** Cross-validation performance evaluation on NetMHCIIpan 4.1 and NetMHCIIpan 4.0. Each dot in the plots refers to a single dataset. AUC 0.1 refers to the area under the ROC curve integrated up to a false positive rate of 10%, and PPV is the predicted positive values calculated from the proportion of positives with the top N highest predictions, where N is the total number of positives within the given dataset [15]. NetMHCIIpan-4.1 significantly outperforms NetMHCIIpan-4.0 in all performance metrics ($p<0.001$, binomial test, in all cases).

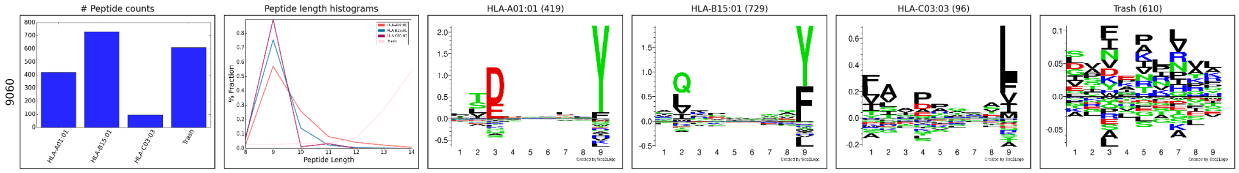


**Supplementary Figure 2. MHCMotifDecon HLA class I motif deconvolution** for the IHW09060 dataset using a trash bin threshold of 2%.

A)

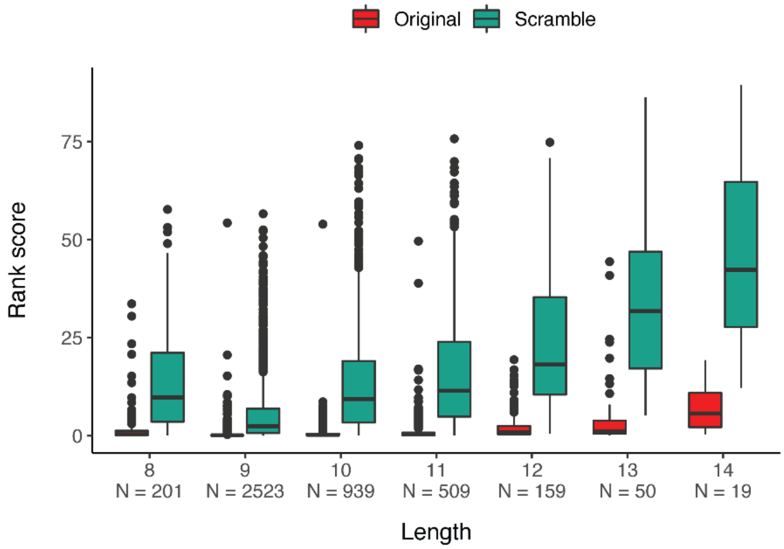


B)

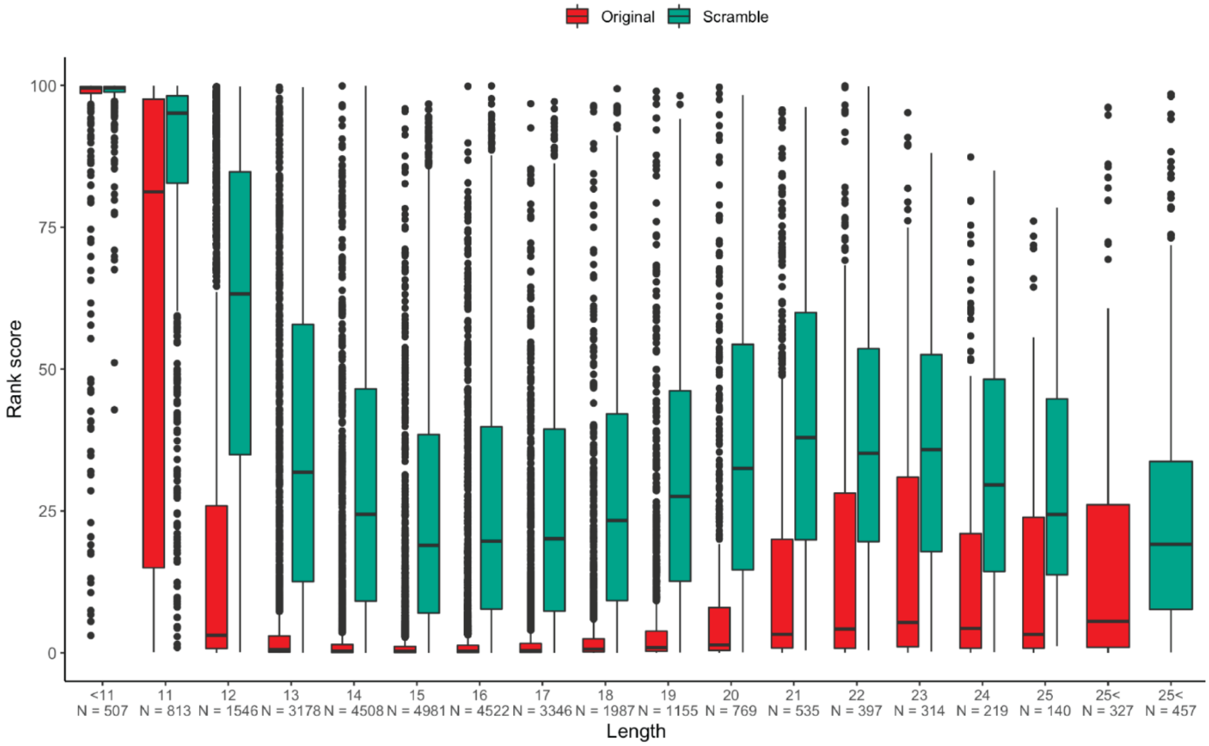


**Supplementary Figure 3. Comparison of rank scores per length after deconvolution of original and scrambled datasets with MHCMotifDecon**. In the scrambled dataset, the peptide sequences have the same amino acid composition but in a shuffled order (for details refer to methods section). 100 scrambled variations were generated for each peptide. For comparison a random sample of all the scrambled peptides matching the number of peptides in the original dataset were selected (for details on scrambled data refer to Methods section). To quantify the proportion of scrambled peptides that would be assigned to an HLA expressed by the cell line, the rank scores obtained by MHCMotifDecon were compared to those obtained from the original peptide data. A) Shows the results for HLA class I, and B) the results for HLA class II.

A)

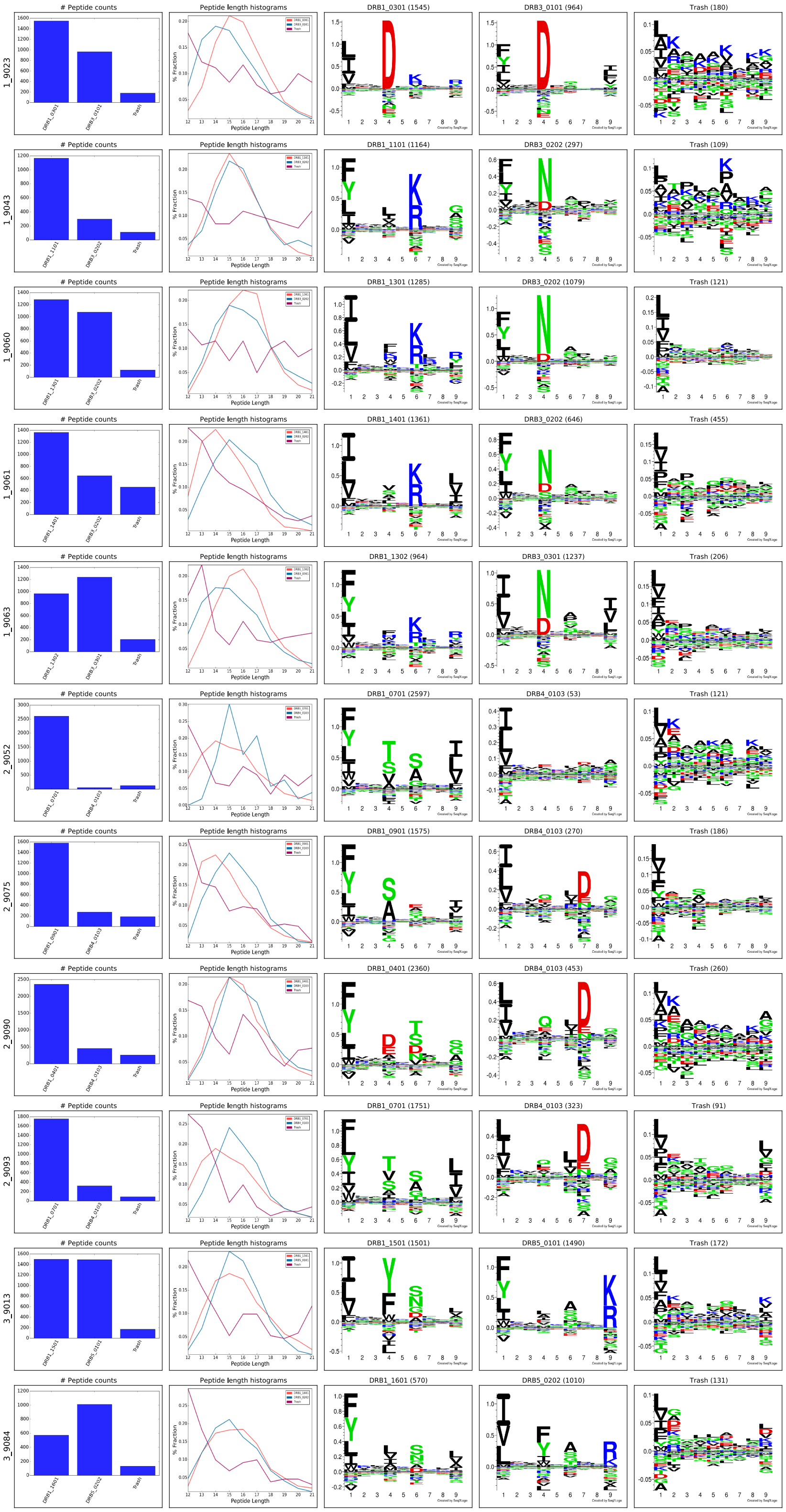
1_9023
### Peptide counts
Peptide length histograms
DRB1_0301 (1545)
DRB3_0101 (964)
Trash (180)
1_9043
DRB1_1101 (1164)
DRB3_0202 (297)
Trash (109)
1_9060
DRB1_1301 (1285)
DRB3_0202 (1079)
Trash (121)
1_9061
DRB1_1401 (1361)
DRB3_0202 (646)
Trash (455)
1_9063
DRB1_1302 (964)
DRB3_0301 (1237)
Trash (206)
2_9052
DRB1_0701 (2597)
DRB4_0103 (53)
Trash (121)
2_9075
DRB1_0901 (1575)
DRB4_0103 (270)
Trash (186)
2_9090
DRB1_0401 (2360)
DRB4_0103 (453)
Trash (260)
2_9093
DRB1_0701 (1751)
DRB4_0103 (323)
Trash (91)
3_9013
DRB1_1501 (1501)
DRB5_0101 (1490)
Trash (172)
3_9084
DRB1_1601 (570)
DRB5_0202 (1010)
Trash (131)
% Fraction
Peptide Length

B)

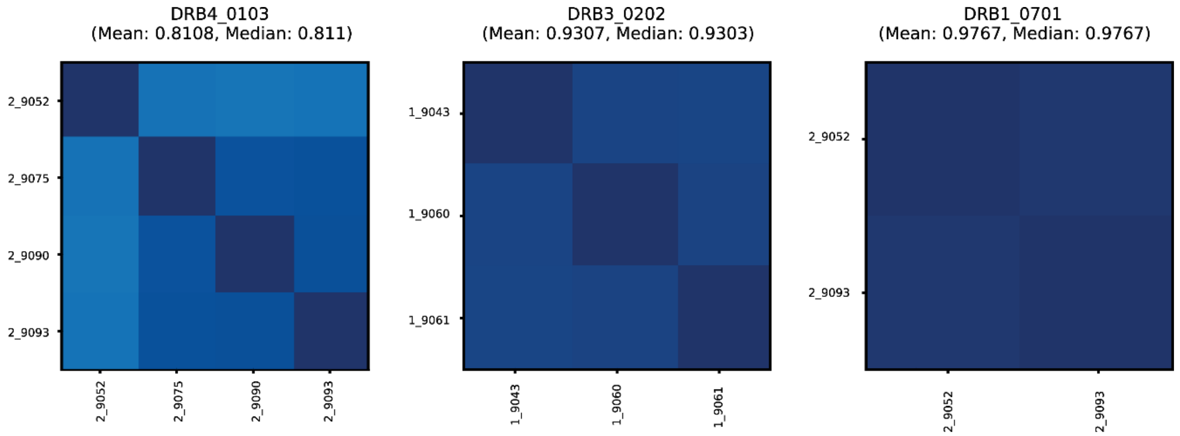


**Supplementary Figure 4. Motif deconvolution of the 11 HLA-DR datasets.** Peptide datasets were filtered for HLA class I binders as described in the text and submitted to MHCMotifDecon for motif deconvolution using default class II options. A) Logo output plot generated by the method. Each row corresponds to one dataset (cell line). The label N_ in front of each cell line (i.e 1_9023) corresponds to the different haplotype groups (see supplementary table 1). The first column shows the number of peptides assigned to each HLA molecule and the Trash bin (containing peptides with a predicted rank > 20%). The second column gives the peptide length distribution for each HLA, and Trash bin, and the remaining columns the binding motifs for each HLA molecule and Trash bin. Motifs are constructed from the predicted binding cores using Seq2Logo [46] with default settings. B) Consistency matrices generated by the method for the three DR molecules shared between 2 or more cell lines using the method described in [15] defining the similarity between two HLA binding motifs in terms of the Pearson's correlation coefficient (PCC) between the two vectors of 9*20 elements (9 positions and 20 amino acid propensity scores at each position). Note. that the consistency plot for DRB4*01:03 includes the null allele DRB4*01:03N expressed in the 9052 cell line. Removing this allele from the plot results in increasing the Mean PCC value to 0.88.
